# Targeted gene sequencing in 6994 individuals with neurodevelopmental disorder with epilepsy

**DOI:** 10.1101/602524

**Authors:** Henrike O. Heyne, Mykyta Artomov, Florian Battke, Claudia Bianchini, Douglas R. Smith, Nora Liebmann, Vasisht Tadigotla, Christine M. Stanley, Dennis Lal, Heidi Rehm, Holger Lerche, Mark J. Daly, Ingo Helbig, Saskia Biskup, Yvonne G. Weber, Johannes R. Lemke

## Abstract

**Purpose:** We aimed to gain insight into frequencies of genetic variants in genes implicated in neurodevelopmental disorder with epilepsy (NDD+E) by investigating large cohorts of patients in a diagnostic setting.

**Methods:** We analyzed variants in NDD+E using epilepsy gene panel sequencing performed between 2013 and 2017 by two large diagnostic companies. We compared variant frequencies in 6,994 panels to other 8,588 recently published panels as well as exome-wide *de novo* variants in 1,942 individuals with NDD+E and 10,937 controls.

**Results:** Genes with highest frequencies of ultra-rare variants in NDD+E comprised *SCN1A, KCNQ2, SCN2A, CDKL5, SCN8A* and *STXBP1*, concordant with the two other epilepsy cohorts we investigated. Only 46% of the analysed 262 dominant and X-linked panel genes contained ultra-rare variants in patients. Among genes with contradictory evidence of association with epilepsy *CACNB4, CLCN2, EFHC1, GABRD, MAGI2* and *SRPX2* showed equal frequencies in cases and controls.

**Conclusion:** We show that improvement of panel design increased diagnostic yield over time, but panels still display genes with low or no diagnostic yield. With our data, we hope to improve current diagnostic NDD+E panel design and provide a resource of ultra-rare variants in individuals with NDD+E to the community.

## Introduction

In recent years, genetic research has gained novel biological insights into the etiology of neurological disorders, particularly in epilepsy^1,2^. Neurodevelopmental disorders with epilepsy (NDD+E) are a rare group of disorders frequently caused by *de novo* events in protein-coding genes^3,4^ precise genetic diagnosis influences genetic counseling but may also guide treatment decisions by enabling medication or treatment tailored to the patient’s underlying genetic defect^2,4^. Examples include treatment with sodium channel blockers in *SCN2A*-and *SCN8A*-related NDD+E^5,6^, ezogabine in *KCNQ2*-related NDD+E^7^ or a ketogenic diet in *SLC2A1*-related GLUT1 deficiency^8^. Up to 28 % of de novo variants (DNV) being found in NDD+E-related genes are associated with such targeted treatment approaches^4^. However, assessments of how often NDD+E-associated genes are mutated are currently insufficient due to lack of large-scale genetic analyses in NDD+E.

Targeted sequencing of specific disease-related gene panels has been part of the diagnostic workup of highly prevalent heterogeneous disorders such as breast cancer^9^, cardiomyopathy^10^ and epilepsy^11-13^. Multiple genes are sequenced in parallel enabling lower sequencing cost, higher coverage and near-absence of secondary findings compared to exome sequencing^14^. However, high heterogeneity of epilepsy gene panel content has been observed^4,15^. This is likely due to the dramatically growing number of genes associated with epilepsy and diverse integration in the established panels, often without robust statistical evidence^4,16^. To increase yield in diagnostic sequencing panels, it is essential considering genes with proven disease association as well as a reasonable frequency of pathogenic variants among affected individuals.

Here, we report likely damaging variants in 645 epilepsy panel genes sequenced at two molecular diagnostic companies, CeGaT (Germany) and Courtagen (USA). In total, 6994 patients with NDD+E of suspected monogenic cause underwent diagnostic sequencing at the respective companies, the majority as first-tier diagnostic test. We compare this large cohort of panels in NDD+E patients with another study of similar design (n=8565)^17^, 10,937 controls as well as with a cohort of exome-wide DNV in NDD+E^4^ (n=1942) investigating variant frequencies in confirmed and putative NDD+E genes in NDD+E panels.

## Materials and Methods

### Gene Panel Sequencing Data

We analyzed gene panel sequencing data of 6994 individuals diagnosed with NDD+E or related disorders of suspected monogenic origin. The data was generated during routine diagnostic sequencing by two different commercial companies, Courtagen (US, n = 3817 cases) and CeGaT (Germany, n = 3177 cases) with similar overall approach and design^11^. Information on cognitive outcome was available in about 59% of cases revealing fractions of individuals with intellectual disability (ID) of 96% (2176/2266, Courtagen) and 97.8% (1833/1875, CeGaT). In the majority of cases, epilepsy was early-onset (before three years of age). Analysis was performed between 2013 and 2017, during which time up to 10 different but vastly overlapping NDD+E panel designs were used by each company. Panels contained a median of 471 and 498 confirmed or suspected epilepsy genes at each respective company and a median 4870 individuals were sequenced per gene (Supplementary Figure S1, Supplementary Table S1, Figure 3). We decided to analyze the 645 genes (see Supplementary Table S2) that were sequenced in at least 2000 individuals. As the first systematic guideline for diagnostic variant interpretation was not published before 2015^18^, we decided to focus on functional (null variants as well as missense variants predicted to be deleterious by in silico tools, see Methods) ultra-rare variants without pathogenicity labels that are not present in the general population^19^. In this setting, functional variants in genes not ordered by the respective clinician were not consistently reported, while we cannot access genes ordered by clinicians. Consequently, we identify few genes with significantly lower variant frequencies in cases than controls (Supplementary Figure S3) suggesting underreporting of variants in these genes.

### Data Processing

A more detailed description of the analysis pipeline of the Courtagen company has been published^20^. A brief overview of analysis steps is described as follows. Courtagen and CeGaT employed custom-designed Agilent Haloplex and SureSelect enrichment kits, to enrich patients’ genomic DNA for target regions of epilepsy (candidate) genes. This was followed by paired-end sequencing (250 or 200bp, respectively) on Illumina platforms (miSeq and HiSeq). Adaptor sequences were then trimmed, and the sequencing reads were aligned to the human reference genome hg19 (GRCh37) with bwa-mem (bio-bwa.sourceforge.net). Reads that mapped equally well to more than one genomic position were discarded. Quality checks were performed ensuring adequate distributions of various quality control metrics such as insert size distribution, mismatch rates, GC bias etc. Subsequent variant calling was done with different pipelines. Variants were filtered for population frequencies < 1% (ExAC, EVS, 1000 Genomes) and platform-specific sequencing artifacts. Follow-up Sanger sequencing was then performed on most variants available to us.

In case of available parental samples, the *de novo* status of individual variants was tested by Sanger sequencing. For one of the companies, segregation testing was partially documented. Out of 1173 ultra-rare damaging variants, 162 (14%) were verified as *de novo*, 36 (3%) segregated with disease and for 975 (83%) segregation was unknown.

### Reannotation and Filtering

All variants reported to patients as well as variants in controls were re-annotated with the following pipeline. Variants were annotated with Ensembl’s Variant Effect Predictor^21^ (=VEP) of version 82 using database 83 of GRCh37 as reference genome. Per variant the transcript with the most severe impact, as predicted by VEP, was selected for further analyses. The decreasing order of variant impacts was HIGH, MODERATE, MODIFIER, LOW. Only protein-altering variants [missense or null (premature stop codon, essential splice site, frameshift)] were included in further analyses. Variants that were present in a subset of ExAC (v0.3), an aggregation of exome sequences from adult individuals without severe childhood-onset diseases and without psychiatric diseases (n = 45,376)^19^, were excluded, as these have been shown to convey no detectable risk to disease on a group level^22^. To increase power for variants that were not tested for segregation, we filtered missense variants predicted to be damaging by PolyPhen^23^ (v2.2.2) or Sift^24^ (v5.2.2). In total, 42% of individuals had no, 34% had one, 15% had two and 8% had three or more ultra-rare variants (either damaging missense or null variant). We labeled ultra-rare variants for which we had no information on segregation as putative *de novo* variants when they had previously been reported as confirmed DNV in individuals with NDD+E^4^ and/or ClinVar^25^ (date 08/2017).

### Population controls

We used controls as a population reference of ultra-rare variant frequencies per gene. The population control dataset was assembled at the Broad Institute from multiple exome sequencing projects. It included data from NHLBI Exome Sequencing Project (for details see http://evs.gs.washington.edu/EVS/), T2D-Genes study (http://www.type2diabetesgenetics.org/projects/t2dGenes), ATVB cohort (dbGAP accession phs000814.v1.p1) and Ottawa Heart study (dbGAP accession phs000806.v1.p1). All control samples were jointly processed through one alignment and variant calling pipeline. Samples of European ancestry were identified using principal component analysis, All 1^st^ degree relatives and duplicated samples were removed from downstream analysis with pairwise IBD analysis in PLINK^26^. From this data, a subset of genes present in diagnostic epilepsy panels was then used as control data in this study, excluding samples with a genotype call rate <95% totaling 10,937 individuals with mean age of 65 with no evidence of psychiatric/neurodegenerative disorder. We subjected genotypes for quality checks keeping only genotypes with > 30X coverage and genotype quality (GQ, estimated in GATK pipeline^27^) > 30. On average, the number of sites with non-reference genotypes in controls that were excluded due to coverage <30X for this analysis was 1.03% (see Supplementary Figure S2). In one company, this number is on average 0.2% (personal communication). Due to the targeted approach, we expect this to be similarly low in the other company. Diagnostic panels may be at an advantage to identifying variants compared to exomes as they have higher average coverage and were subjected to initially lower GQ cutoffs. On the other hand, variants have been validated by Sanger sequencing in some of the controls, but all of the cases and variants in certain genes were systematically underreported in panels (see Figure 1). Controls are of non-Finnish European origin while cases are mostly of non-Finnish European origin, with few exceptions (personal communication). While controls and cases were not matched for more specific population structure we expect this to have no significant influence in singleton rates as these are relatively consistent in different (particularly non-Finnish European) populations in the 1000 genomes project (https://www.nature.com/articles/nature15393/figures/1) >and we also show, that many singletons in cases are likely of *de novo* origin.

**Figure 1.**
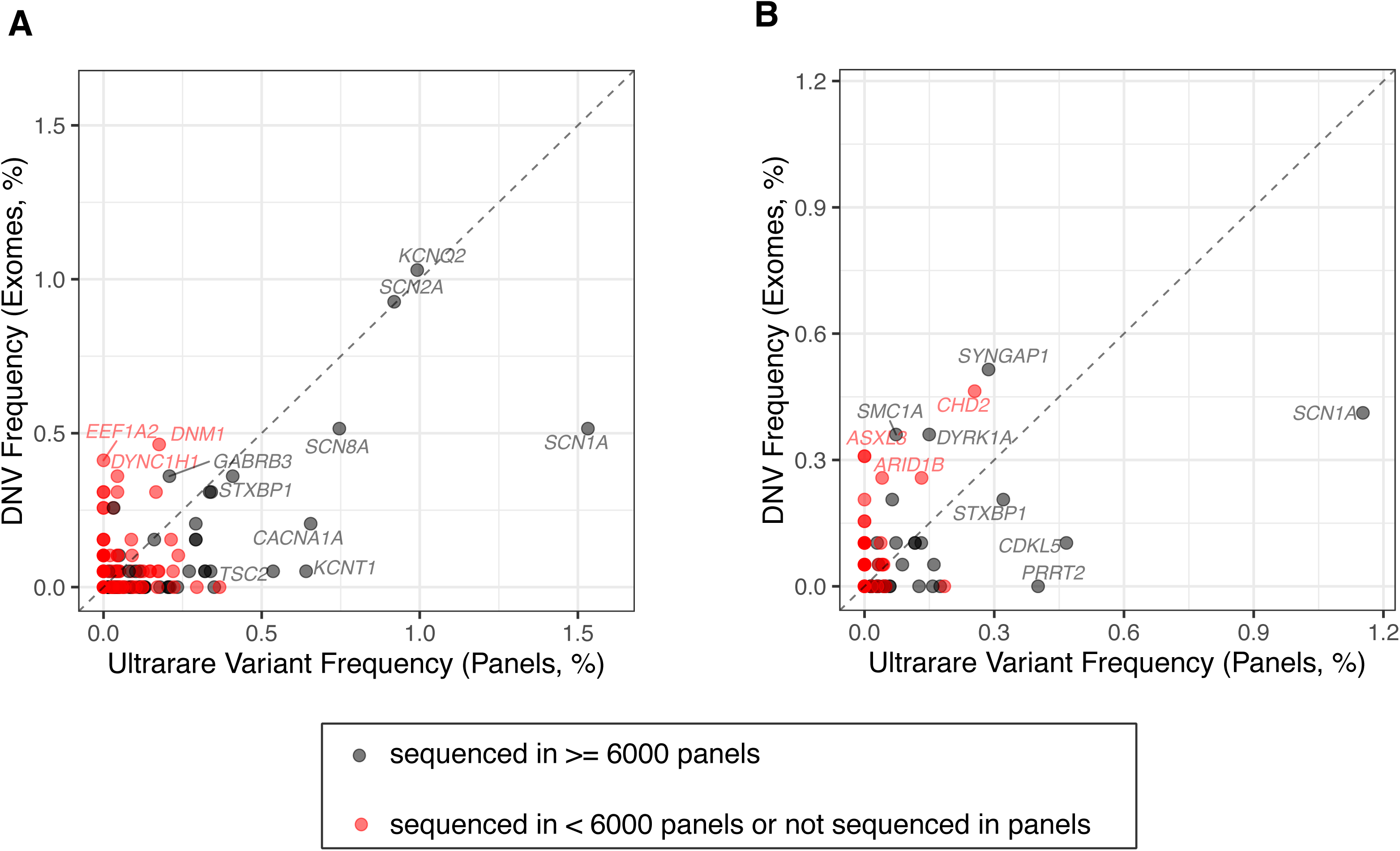
Ultra-rare variants in panels compared to DNV in exomes. **A)** damaging missense variants; **B)** null variants. Genes that were sequenced in >= 6000 panels are labelled black, else red. The dotted line represents equal frequency of *de novo* variants in exomes and ultra-rare variants in panels. Only variants in confirmed disease genes are shown (see methods). DNV frequency in *SCN1A* should be depleted as it is only occasionally pre-screened prior to panel, but usually pre-screened prior to exome sequencing. Frequencies of DNV in exomes and ultra-rare variants in panels are correlated when considering highly covered panel genes (black dots). Missense variants: p-value = 3×10^-9^, rho= 0.63; null variants: p-value= 4×10^-6^, rho= 0.53, method: Spearman correlation.

### Statistical analyses

We assessed individual gene tolerance to null or missense variants in the general population by using the pLI score (probability of being loss-of-function intolerant), missense z-score (z-score of observed versus expected missense variants)^19^ or s_het_ score (selective effects for heterozygous protein null variants)^28^. We defined a gene as constrained with the cut-offs > 0.9 for pLI, > 3.09 for missense z-scores and > 0.05 for s_het_ based on recommendations of the score developers. We compared scores using Wilcoxon rank tests pLI and s_het_ scores as the data appeared not normally distributed upon inspection. As disease gene reference, we used a curated list of disease genes compiled by clinicians as part of the DDD study (http://www.ebi.ac.uk/gene2phenotype/downloads/DDG2P.csv.gz, version 11/7/2018). We subset the list to genes associated with any descending HPO terms^29^ of epilepsy (HP:0001250) or intellectual disability (HP:0001250) or “Brain/Cognition” and only included dominant/X-linked disease genes labelled as “confirmed” or “probable”. We also annotated MPC scores (for Missense badness, PolyPhen-2, and Constraint), a pathogenicity score that leverages regional depletion of missense variants in the general population as well as amino acid deleteriousness^30^ to compare ultra-rare and DNV.

### Code availability

All statistical analyses were done with the R programming language (www.r-project.org). The code will be available upon request.

## Results

### Genes with ultra-rare variants in NDD+E include DEE but also NDD genes

We assessed frequencies of likely protein-altering (missense or null) ultra-rare variants in 6,994 individuals with NDD+E (Figure 1). While we did not assess variant pathogenicity with all ACMG criteria^18^, this class of variants should be enriched for likely pathogenic variants. We analyzed 645 genes that were sequenced in at least 2000 individuals with NDD+E, with a median of 4870 individuals sequenced per gene. Of these, 215 genes were annotated as acting in an autosomal dominant, 47 X-linked, 329 autosomal recessive and 54 unknown inheritance mode. It has been shown repeatedly^22,31^, that genes contributing to severe childhood-onset diseases with high penetrance are depleted for missense/null variants in the general population, measured by pLI/missense z-score^19^. Genes classified as constrained by a significant pLI/missense z-score likely contribute to NDD+E in a dominant/X-linked mode. Of 262 dominant/X-linked genes, 85 genes were constrained and carried at least two ultra-rare variants in our dataset. 41 of these 85 genes were previously described as developmental and/or epileptic encephalopathy (DEE/EE) and NDD+E genes^4,32,33^, while other frequently mutated genes were associated with other well-known genetic syndromes (e. g. *BRAF, KMT2D, TCF4*) or structural brain abnormalities (e. g. *ARX, CASK, FLNA, TUBB4A*). We compared per-gene ultra-rare variant frequencies to 10,937 controls assessing the general population background rate (Supplementary Figure S3). Ultra-rare variant frequencies (missense and null) of the top genes were *SCN1A* (2.7%), *KCNQ2* (1.2%), *SCN2A* (1.0%), *CDKL5* (0.8%), *SCN8A* (0.8%) *STXBP1* (0.7%) and *CACNA1A* (0.7%). Reassuringly, ranks of top genes were in concordance with a recently published study of similar design (gene panel sequencing in 8565 epilepsy patients^17^, see Figure 2).

**Figure 2.**
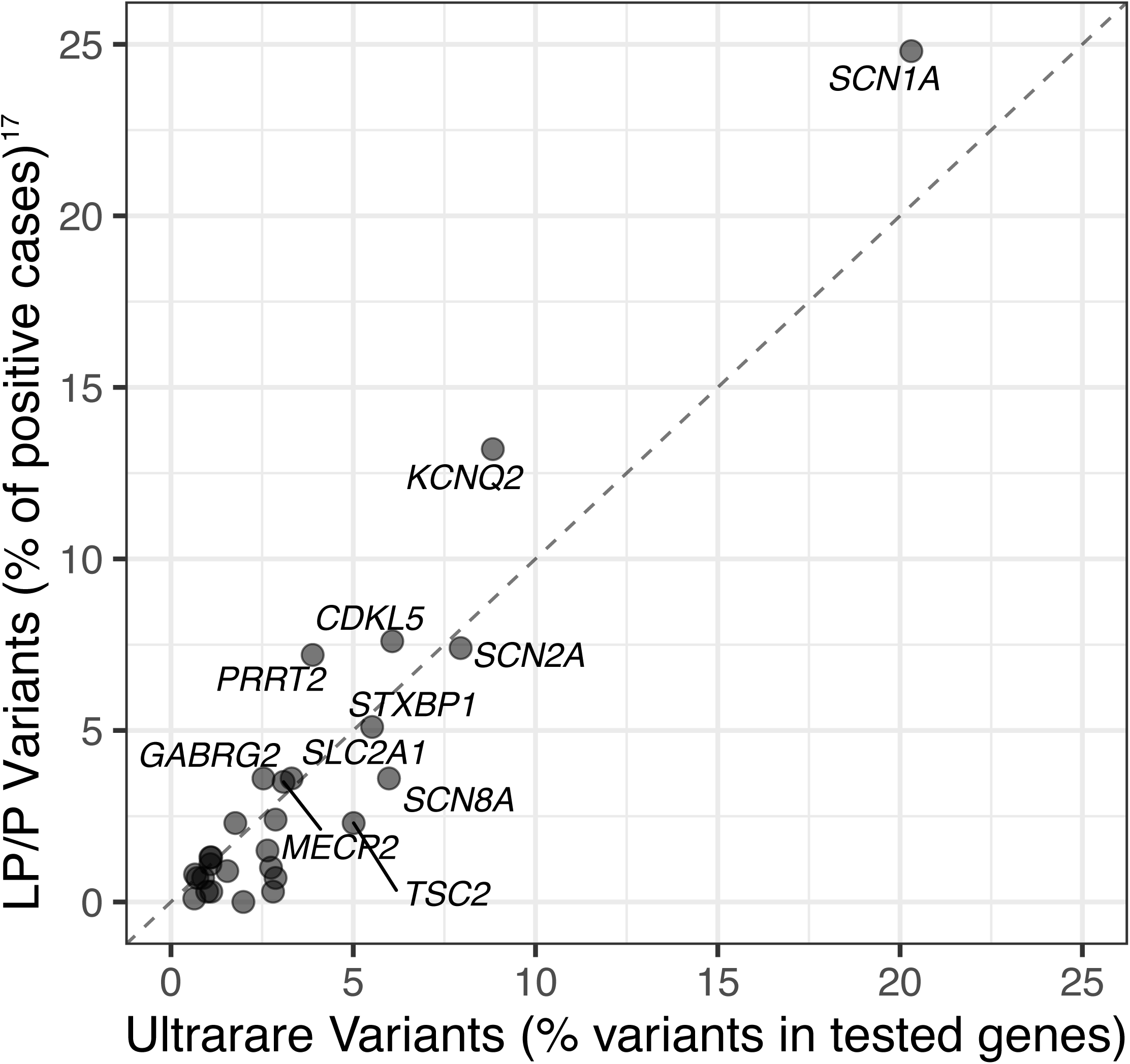
Pathogenic variants in an independent panel cohort. Ultra-rare variants in panels (damaging missense + null) versus pathogenic variants in a panel cohort of 8585 individuals including damaging missense + null + CNV (CNVs constitute about 9% of pathogenic variants)^17^. Adapted to the format of^17^, the fraction of pathogenic variants in each gene is given as the proportion of variants in all positive cases. Only genes included in Lindy *et al.* ^17^ are shown. Correlation of data shown: p-value= 4×10^-7^, rho= 0.79, method: Spearman correlation.

**Figure 3.**
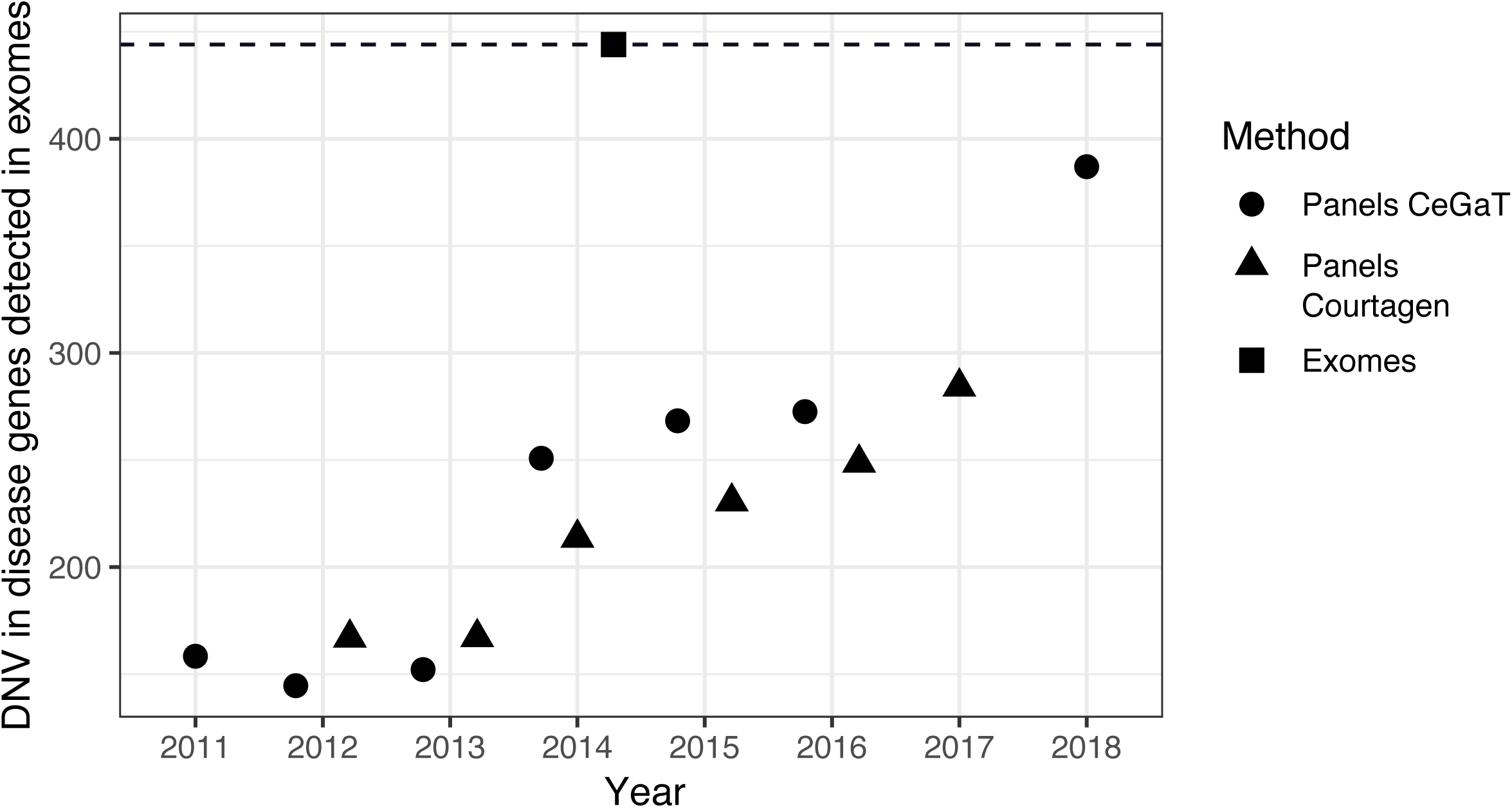
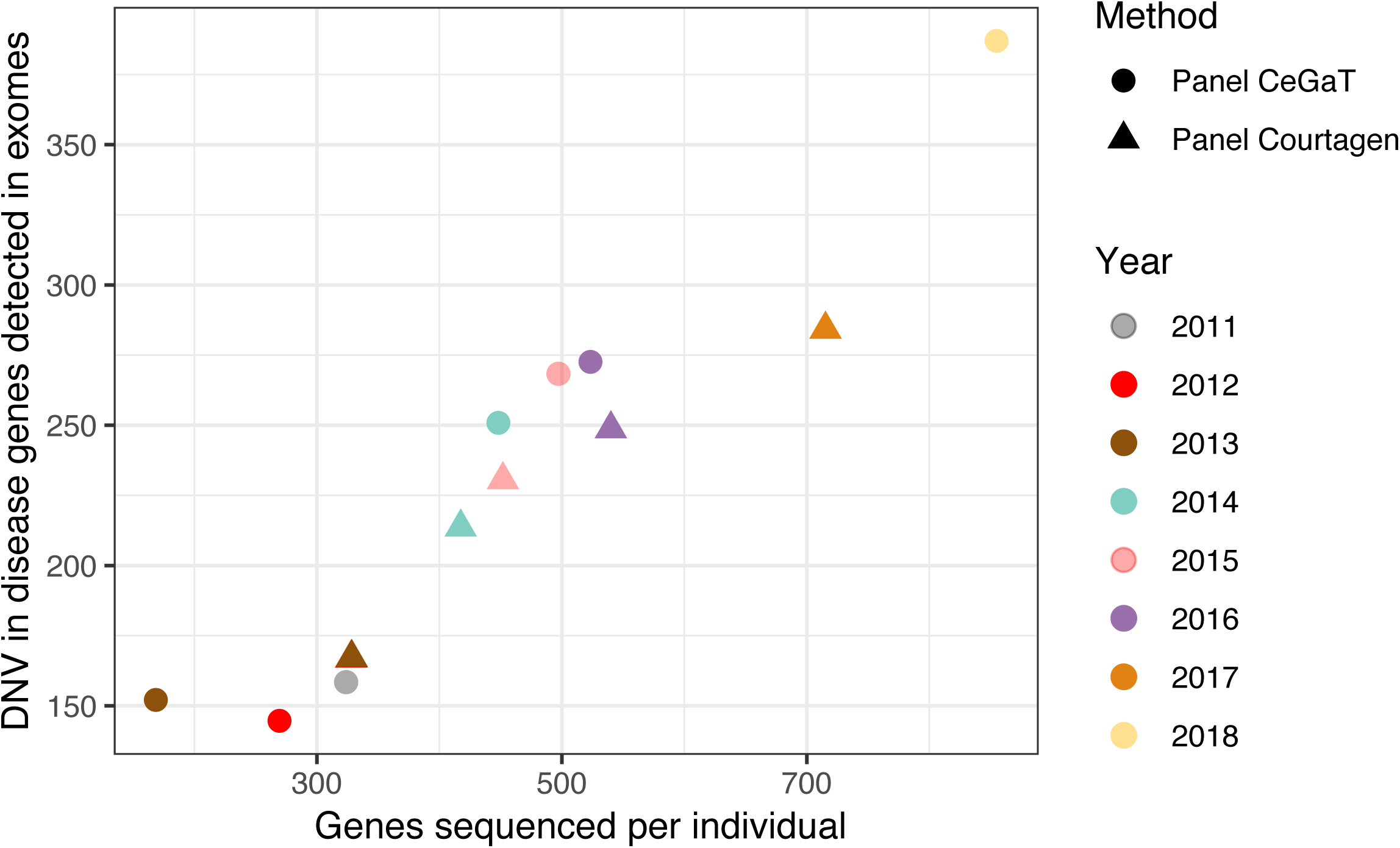
Number of disease genes in panels increasing over time. We determined how many DNV in disease genes (as reported by the Deciphering Developmental Delay study^31^, see methods) would have been found in 1942 individuals with NDD+E as part of an exome-wide study^4^ when using panels instead of exomes. For example, of the 444 exome-wide DNV detected in Heyne et al. 2018^4^, the panel designs of our current study of the years 2011 to 2013 would have covered less than 200. After improvement of panel design over the years, up to 300-400 DNV would have been detected in 2017 to 2018. The increasing number of DNV in disease genes is correlating with increasing panel sizes over time. As we do not evaluate individual variant pathogenicity and do not include all disease genes, true diagnostic yields would be different and likely higher (usually up to 40% for exomes^34,35^ and up to 26% for panels in 2018). **A)** Number of individuals with DNV in disease genes in panels over time (triangles and dots) versus exomes (square). **B)** total number of genes sequenced per patient and number of diagnoses

### Comparing variant frequencies per gene in 6,994 panels and 1,942 trio exomes

We compared ultra-rare variant frequencies in our panel dataset to DNV frequencies in a large recent exome-wide trio study of 1,942 individuals with NDD+E^4^. Restricting our dataset to genes sequenced in 6000 to 6994 individuals we found correlation between the datasets for both missense and null variants (Missense variants: p-value = 3×10^-9^, rho= 0.63; null variants: p-value= 4 ×10^-6^, rho= 0.53, method: Spearman correlation, see Figure 1). That suggests, that a large fraction of ultra-rare variants in our dataset arose *de novo* even if only a fraction of them were tested for segregation. However, there was no or negative correlation (Missense variants: p-value = 0.05, rho= −0.17; null variants: p-value= 0.7, rho= −0.04, method: Spearman correlation) between panels and exome sequencing when considering all genes, as many genes were not included in the diagnostic gene panels. Assuming gene panel sequencing identifies 100% of the DNV found in trio exome sequencing for a given gene, we investigated how many likely protein altering DNV in curated disease genes (see methods) would have been found in 1942 individuals with NDD with epilepsy as part of the exome-wide study when using panels instead of exomes. We found 444 DNV in the exome sequencing data in those genes, while panels would have identified on average 245 DNV. The proportion of identified DNV in panels significantly increased over time though, as panels were continuously updated according to the literature (Figure 3). In this approach we only consider damaging missense and null DNV and do not evaluate pathogenicity of individual variants. Therefore and as the set of disease genes is more strictly defined, true diagnostic yields are likely higher (usually up to 40% for clinical exome sequencing^34,35^ and up to 26% in most recent panel diagnostics).

### The majority of genes contained no or fewer ultra-rare variants in epilepsy cases than in controls

Comparing variants in cases and controls, we noticed that 255 out of 645 panel genes (39.5%) did not display any ultra-rare variants in > 2000 NDD+E cases (Figure 1, Supplementary Figure S3, Supplementary Table S2). Further 247 genes (38.3%) had lower frequencies of ultra-rare variants in cases compared to population controls. The majority of these in total 502 rarely mutated genes were of autosomal recessive inheritance (60%, 300/502), for which we would not expect higher variant frequencies in cases. However, 30% (149/502 genes) were of autosomal dominant/X-linked inheritance (Supplementary Table S2). For the remaining 10%, inheritance was unclear. A limitation of this study is that we cannot guarantee, that variants in genes not ordered by clinicians were consistently reported and therefore we cannot exclude that some missed variants were in true disease genes. However, the 149 not or rarely mutated dominant/X-linked genes had lower constraint scores (pLI: 0.93 [median], missense z-score: 2.2±2.6 [mean, SD], s_het_ score: 0.08 [median]) on a group level than the 119 dominant/X-linked genes with higher ultra-rare variant frequencies (pLI: 0.99 [median], missense z-score: 3.5±2.1 [mean, SD], s_het_ score: 0.16 [median]) in cases than in controls (see Methods, respective p-values pLI score: 5×10^-4^, missense z-score: 2×10^-5^, s_het_ score: 1×10^-4^). Additionally, they were not significantly different in estimated mutation rate^36^ (two-sided t-test, p-values for missense mutation rate 0.76, null mutation rate 0.86). This suggests that a lower mutation rate is not the reason for low frequencies of ultra-rare variants in most of these genes among NDD+E cases, instead it is more likely that most of these genes are not true NDD+E genes.

### Confirmed and putative de novo variants

Among 6994 epilepsy cases, we revealed 333 DNV that were not in ExAC as well as either damaging missense or null DNV. 95% (317/333) of DNV were in 54 constrained genes, 32 genes displayed at least two DNV (Supplementary Table S1, Figure 1, Supplementary Figure S4). 4.7% (331/6994) of cases had a total number of 333 damaging DNV. As segregation testing was not performed systematically in the overall cohort, this number is certainly an underestimate (see Methods). It has been documented for many disease genes including genes associated with NDD+E^37^, that disease-causing missense variants cluster in particular functionally relevant protein domains. We annotated MPC scores, a pathogenicity score that considers if missense variants in the general population are depleted in particular regions of a gene^30^. Higher MPC scores indicate increased deleteriousness of missense variants. We found a median MPC score of 2.3 for 333 DNV and 0.76 for 11,233 ultra-rare variants for which disease segregation was unknown (Wilcoxon-rank sum test, p-value 1×10^-76^). Also within constrained genes, we found a median MPC of 2.13 in DNV and 1.03 for variants with unknown segregation status (Wilcoxon-rank sum test, p-value 2×10^-45^). These results confirm the increased likelihood of pathogenicity of DNV in comparison to ultra-rare variants with unknown disease segregation.

## Discussion

Gene panel analysis is widely used in genetic diagnostics of NDD+E. However, panel designs vary substantially across companies^4,15^ and over time. Here, we report a large cohort of individuals with NDD+E (6,994 cases) that underwent gene panel sequencing in a diagnostic setting.

Frequencies of ultra-rare variants in our cohort were compared with two other large NDD+E cohorts: 1) DNV in 1942 trio exomes^4^ and 2) likely pathogenic variants in 8565 gene panels^17^. Of the top 20 disease genes with the highest numbers of DNV in exomes, 16 were also present in our panel data. *ARID1B, ASXL3, EEF1A2* and *SLC6A1* were the genes missing in panels. Considering the top 35 disease genes in exomes (at least 4 DNV in exomes) missing in panels were also *KCNH1, PURA, COL4A3BP KIF1A, ANKRD11, DDX3X, MED13L* and *PPP2R5D*. We suggest those genes could be added in subsequent panel designs. Of the top 20 exome genes, only 8 were present in Lindy et al^17^. These results illustrate the high genetic heterogeneity of NDD+E. The most frequently mutated genes in exomes as well as panels were *SCN1A, SCN2A* and *KCNQ2*. Following at about half their frequency were *CDKL5, SCN8A, STXBP1, SYNGAP1, TSC2* and *CACNA1A.* These genes are consistently present at higher diagnostic yield in NDD+E^11,13,34^. Of note, *GABRG2, TSC2* and *PRRT2* had high frequencies of ultra-rare variants in our panel study and in Lindy *et al*^17^ but barely displayed DNV in exomes (*GABRG2* and *TSC2*: 1 DNV, *PRRT*: 0 DNV) suggesting that many of the variants in *TSC2, GABRG2* and *PRRT2* may be inherited rather than de novo.

While many disease genes affected in trio exomes were not included in panel designs, we show that gene panel content consistently improves over time. Many frequently mutated genes are associated with “classic” developmental and/or epileptic encephalopathies, whereas others are associated with more unspecific diagnoses of NDD. A too narrow target on “classic epilepsy genes” therefore neglects, that NDD are accompanied by epilepsy in ca. 20% of cases and therefore any NDD gene is principally suggestive to be associated with epilepsy^4,31^. Aptly, we recently showed that 24 diagnostic providers of panel sequencing also lacked a substantial fraction of NDD+E-associated genes in their panel designs^4^. In the early days of NGS, small panel sequencing allowed the introduction of this new technology into clinical diagnostics. Today, panels still offer a cost-effective method to diagnose causal pathogenic variants in the most commonly affected genes as in some countries current reimbursement frameworks do not adequately cover the additional costs of exome sequencing. Yet exome sequencing covers far more of the genetic heterogeneity of NDD+E and a recent American study found that panels are not necessarily more cost-effective than exome sequencing in the US^38^. The number of NDD+E disease genes is continuously increasing, which has only become possible by wide adoption of in particular trio exome/genome sequencing approaches. Detection rates with panel diagnostics are necessarily limited by medical knowledge at the time of panel design. On the other hand, higher coverage in panels than in exomes is superior in detecting low-grade mosaicism in a patient.

The majority of dominant/X-linked panel genes (502 of 645) did either not display any ultra-rare variants in > 2000 epilepsy cases or even had lower frequencies of ultra-rare variants in cases than controls. This could be due to a low mutation rate of these genes or a phenotype rarely ascertained in our cohort. However, given the fact that these genes had no significantly different mutation rate but significantly lower constraint scores compared to all other dominant or X-linked genes in this study it is likely that many of them are not disease associated. This observation is paralleled by a study of similar design on 7,855 individuals with childhood-onset cardiomyopathy, where several genes frequently sequenced in clinical routine, could also not be convincingly associated with disease^10^. In our study, panel design originated in 2010, when multiple candidate genes for rare diseases were nominated without sufficient statistical evidence and could not be confirmed in a clinical setting^39^. This was also described specifically for epilepsy genetics^4,16^.

Of 645 panel genes in our study, 329 genes were associated with recessive inheritance. However, variants in recessive genes segregating with disease were only observed in approx. 1.7% (27 out of 1633 cases) within a documented sub-fraction of this study. This is in concordance with rates of 1.3%^12^ (n = 775 cases) and 1.1%^17^ (n = 8565 cases) in two recent NDD+E studies using gene panels and 3.6% (n = 7,448 cases) in an exome-wide study on developmental disorders with and without epilepsy from non-consanguineous families^40^. Thus, panel designs usually display an imbalanced distribution of recessive genes (very few percent of diagnoses but approximately half of panel genes) versus dominant genes (vast majority of diagnoses but only half of panel genes).

Limitations of our study include inconsistent variant reporting in cases and that the different cohorts we are comparing were neither technically nor ancestry matched. However, we do not expect these technical limitations to alter the key conclusions of this study (see Methods).

We also evaluated the frequencies of ultra-rare variants in five genes with contradictory evidence of gene-disease relationship, which thus had been classified as “disputed” by the formal criteria of the ClinGen Consortium^16^ (*CACNA1H, CACNB4, EFHC1, MAGI2, SRPX2*) as well as two genes with contradictory susceptibility to epilepsy (*CLCN2, GABRD*). *CACNB4, EFHC1, MAGI2, SRPX2, CLCN2* and *GABRD* showed identical frequencies of ultra-rare variants in cases compared to controls (Supplementary Figure S5). Thus, our findings support the evidence that *CACNB4, EFHC1, MAGI2, SRPX2, CLCN2* and *GABRD* may not be truly associated with epilepsy. Coverage of *CACNA1H* was too poor in controls from ExAC to allow a valid comparison of variant frequencies between cases and controls.

In summary, our data provides evidence to further improve the design of NDD+E panels by i) including genes with highest burden of ultra-rare variants, ii) adjusting the ratio of autosomal dominant and X-linked genes with high diagnostic yield versus autosomal recessive genes with low diagnostic yield and iii) excluding genes with poor evidence for true disease-association or very few ultra-rare variants in epilepsy cases.

**Table 1.** – Cohort description

| **Cohort** | **Phenotype** | **pmid** | **n** | **Type of variants** | **Number of genes analyzed** |
| --- | --- | --- | --- | --- | --- |
| CeGaT | NDD+E | unpublished | 3,177 | ultra-rare, DNV | 645 |
| Courtagen | NDD+E | unpublished | 3,817 | ultra-rare, DNV | 645 |
| Lindy *et al*<sup>17</sup><br>(GeneDx) | NDD+E | 29655203 | 8,565 | likely pathogenic<br>(ACMG) | 70 |
| Heyne *et al*<sup>4</sup> | NDD+E | 29942082 | 1,942 | DNV | 18,228 |
| exomes from<br>different<br>cohorts,<br>see Methods | controls | see Methods | 10,937 | ultra-rare | 645 |

NDD+E – neurodevelopmental disorders with epilepsy
DNV – *de novo* variants
ACMG – variant interpretation guidelines by the American College of Medical Genetics

## Acknowledgements

We would like to thank the team of the Institute of Human Genetics of the University of Leipzig as well as the Analytic and Translational Genetics Unit, Boston, for helpful discussions. HOH was supported by stipends from the Federal Ministry of Education and Research (BMBF), Germany, FKZ: 01EO1501 and the German Research Foundation (DFG): HE7987/1-1 and HE7987/1-2. YGW was supported by DFG grants WE4896/3-1 and WE4896/4-1.

## Author contributions

HOH wrote and FB substantially revised the manuscript. HOH, MA, CB performed analyses and made figures. Study design: JRL, SB, DRS, HOH. Data preparation: FB, DRS, CMS, VT, YW, NL, MA, HOH, SB. Interpretation: HOH, JRL, MJD, YGW, HR, FB, IH, HL, DL. All authors approved the final manuscript.

