## Supplemental Information for "Targeted gene sequencing in 6994 individuals with neurodevelopmental disorder with epilepsy"

### SUPPLEMENTARY INFORMATION

#### Supplementary Figures

**Supplementary Figure S1**. **Number of individuals sequenced per gene (histogram)**. Due to differing gene content of individual gene panels the number of individuals sequenced differs per gene (median 4870 individuals, interquartile range 2803 to 6202 individuals).

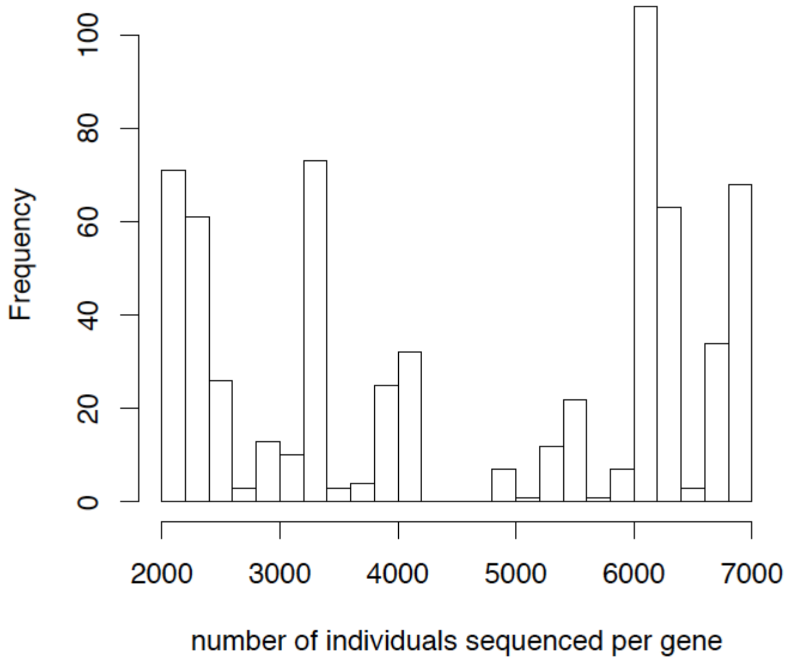

**Supplementary Figure S2**. Distribution of coverage in controls (density).

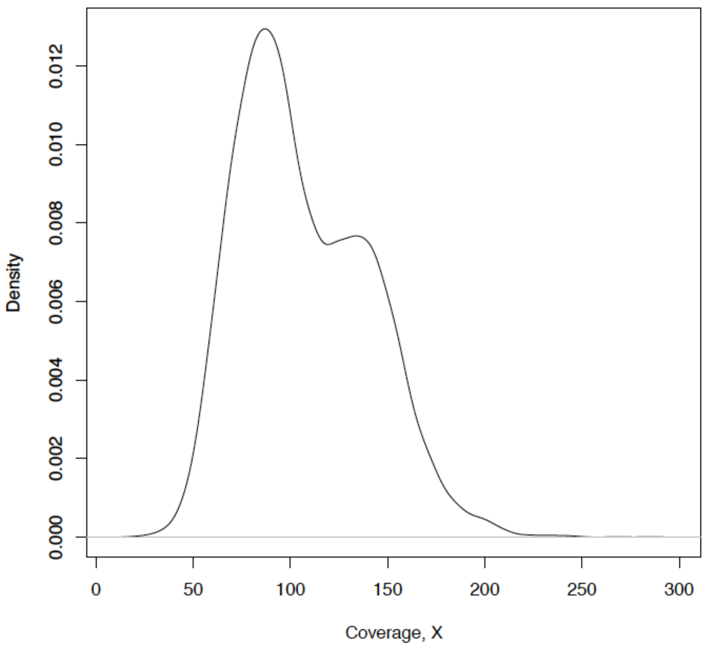

**Supplementary Figure S3. Frequency of ultra-rare variants in cases compared to controls.** Selected genes are labeled. In cases, we only have access to variants that were reported to patients. Therefore, in some clearly not disease associated genes such as *FLG* and *DNAH7* (labelled) variants are likely underreported, resulting in these genes with seemingly higher variant frequencies in controls. **A**, damaging missense variants **B**, null variants.

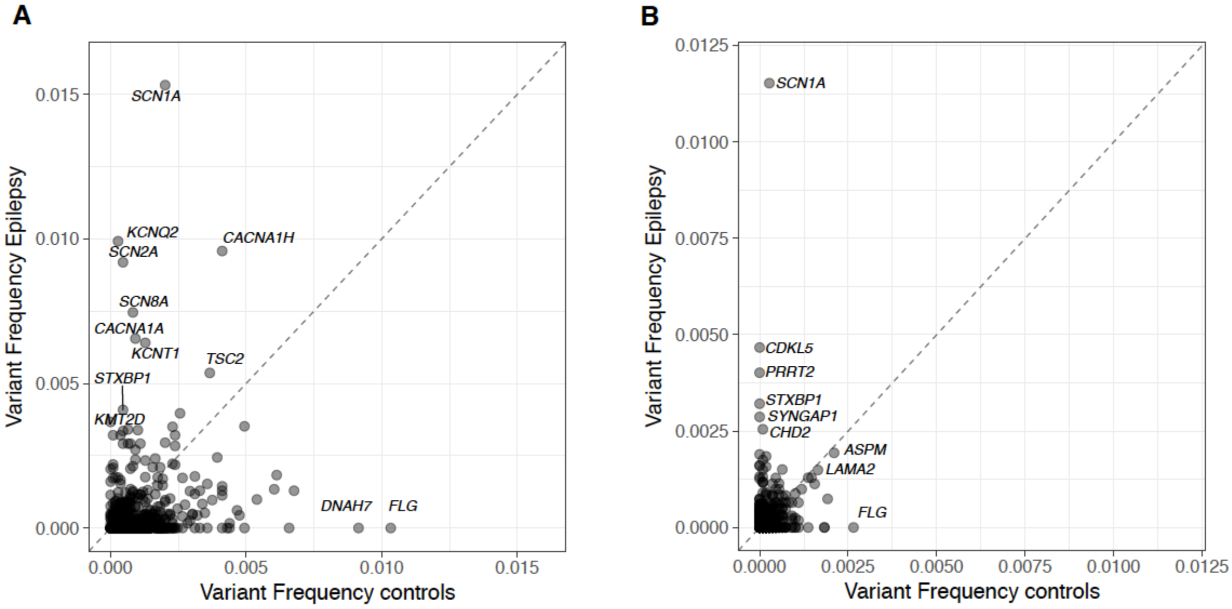

**Supplementary Figure S4. *De novo* variants normalized to number of sequenced individuals in all genes with at least two DNV.** Missense DNV predicted to be damaging (=”dmis”) are plotted in orange, while null DNV are plotted in purple. Variants for which no information on segregation was available, but that have previously been reported as DNV in ClinVar were annotated as putative DNV and colored in light purple or orange, respectively.

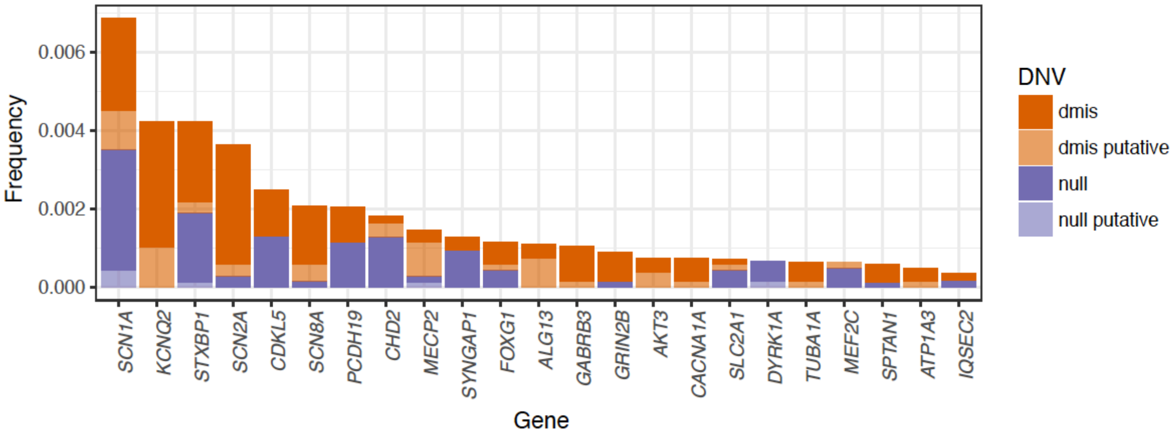

**Supplementary Figure S5. Frequency of ultra-rare variants in cases compared to controls** in five genes with contradictory evidence of association with epilepsy, which had been classified as “disputed” by the formal criteria of the ClinGen Consortium[1]. Correlation: rho = 0.94, p-value = 0.02, method: Spearman correlation.

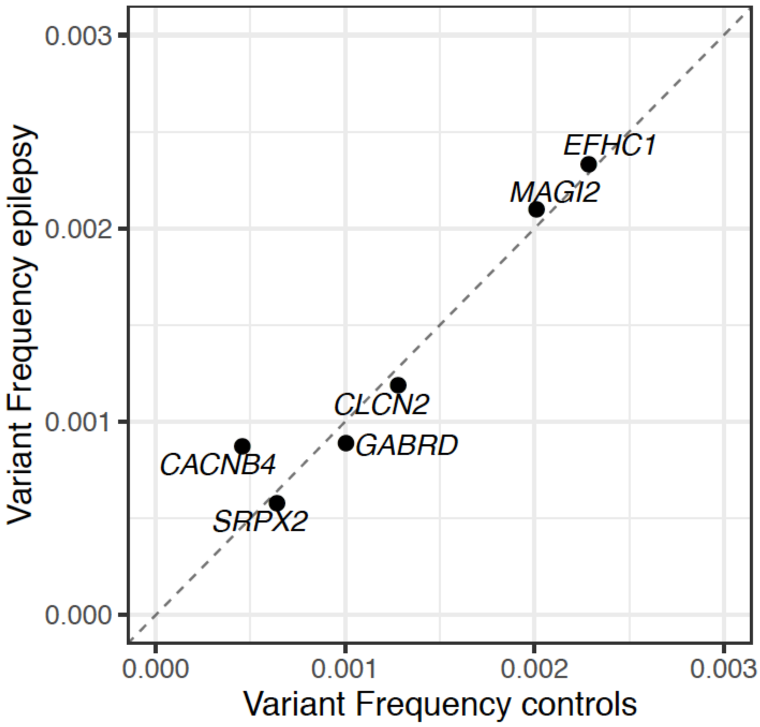

### Supplementary Tables

**Supplementary Table S1**: In this table each line contributes to a variant found in an individual with NDD+E. Variants were annotated with Variant Effect Predictor (please see methods and [2] for details). Additional column names include segregation, MPC[3] and whether variants have previously been reported as *de novo* variants ("DNV_pub", "dnv" for combination of previously reported and confirmed *de novo* variants).

**Supplementary Table S2**: In this table each line contributes to one of 645 genes that were sequenced on different panels in at least 2000 individuals with NDD+E. Columns include number of individuals sequenced per gene (="nseq"), constraint scores pLI, missense z- (="misz) and $s_{het}$ score (see methods), infant brain gene expression (="infant_brain_RPKM") see Heyne *et al*[4], whether genes are annotated as known epilepsy genes (see results), variant counts of null/truncating ("=trunc") and damaging missense ("=dmis") variants and frequencies (="freq") in epilepsy cases and controls.
